# TW-GMProt: Trust-Weighted Semi-Supervised Gated Modulation Protein Language Model for Antimicrobial Peptide Activity Prediction

**DOI:** 10.64898/2026.11.22.768079

**Authors:** Prem Singh Bist, Jagath C. Rajapakse

## Abstract

Antibiotic resistance poses a major global health threat, underscoring the need for faster approaches to antimicrobial discovery. Antimicrobial peptides (AMPs) are promising therapeutic candidates, but experimental activity screening remains costly and time-consuming. We introduce TW-GMProt, a trust-weighted semi-supervised gated modulation protein language model for quantitative AMP activity prediction. It integrates pretrained protein language model representations with convolutional neural networks through a gated modulation strategy, complemented by biophysical priors. To address the scarcity of experimental data, the framework further employs trust-weighted semi-supervised learning to leverage a large collection of unlabeled antimicrobial peptides through pseudo-labeling. For minimum inhibitory concentration (MIC) prediction against Escherichia coli, TW-GMProt achieved an RMSE of 0.434, Kendall’s *τ* of 0.582, *R*^2^ of 0.622, and Pearson’s *r* of 0.792. Compared with MSCMamba, these results correspond to a significant reduction of over 17% lower RMSE (*p* = 1.95 *×* 10^*−*3^), over 25% higher Kendall’s *τ*, over 30% higher *R*^2^, and more than 15% higher Pearson correlation. Comparable predictive performance was observed for Staphylococcus aureus, supporting the applicability of the framework across Gram-negative and Gram-positive bacteria. These results demonstrate that combining gated multimodal representation learning with trust-weighted semi-supervised training provides an effective strategy for AMP activity prediction.

## Introduction

Conventional antibiotic treatments are becoming less effective due to the rapid emergence of resistant pathogens and are often associated with adverse side effects. This growing challenge underscores the urgent need for alternative therapeutic strategies (Behera et al., 2025). Antimicrobial peptides (AMPs) have emerged as promising candidates because of their broad-spectrum activity against bacteria, viruses, and fungi, as well as their potential for targeted action (Alzain et al., 2025; Min et al., 2024; Bhattarai et al., 2025). AMPs are short amino acid sequences, typically 2–100 residues long, that can exert antimicrobial effects through mechanisms including disruption of microbial membranes and interactions with intracellular targets (Zhang et al., 2021).

Most existing computational AMP studies focus on qualitative classification, determining whether a peptide exhibits antimicrobial activity (Xing et al., 2023; Chen et al., 2025; Wang et al., 2024). However, therapeutic development also requires quantitative assessment of peptide potency, particularly the prediction of minimum inhibitory concentration (MIC) values and the reliable ranking of candidate peptides for experimental validation. Accurate activity regression is therefore essential for distinguishing peptides with different levels of antimicrobial efficacy. MSCMamba (He et al., 2025) addressed this problem by reformulating several established AMP classification architectures as regression models and introducing a Mamba-based framework for quantitative activity prediction. Although this approach improved performance over multiple baselines, the limited availability of experimentally measured MIC values remains a major obstacle to learning robust and generalizable activity representations.

In this work, we propose *TW-GMProt*, a semi-supervised, trust-weighted gated protein language modeling framework for quantitative antimicrobial peptide activity prediction. At its core, the model integrates pretrained ProtT5 representations with CNN-guided gated modulation to combine global contextual information with complementary local sequence patterns, while physicochemical, BLOSUM62, and positional descriptors provide additional biochemical, evolutionary, and sequence-order information. To address the scarcity of experimentally measured MIC values, the model extends this architecture through trust-weighted pseudo-label learning. A teacher model generates pseudo-MIC labels for a substantially larger collection of AMP sequences, which are incorporated into training with reduced weights to reflect their lower reliability relative to experimentally measured MIC values. Experimental labels therefore remain the dominant source of supervision during joint training. The resulting model is subsequently fine-tuned using experimentally labeled data only, enabling model to exploit a broader peptide sequence space while limiting the influence of potentially inaccurate pseudo-labels.

TW-GMProt therefore combines gated multimodal representation learning with controlled semi-supervised supervision in a unified framework for MIC prediction. Experimental evaluation against state-of-the-art methods demonstrates improved predictive accuracy and peptide ranking performance, while statistical analysis confirms that the trust-weighted training strategy provides a significant benefit beyond the original model. By leveraging experimentally validated MIC measurements together with large-scale pseudo-labeled peptide data, the model provides a practical approach for improving AMP activity prediction and prioritizing promising candidates for experimental validation.

## Materials and Methods

### Datasets

The primary experimentally labeled minimum inhibitory concentration (MIC) dataset used in this study was obtained from the GRAMPA database (Filippov and Shkirskaya, 2019), an integrated resource that consolidates antimicrobial peptide (AMP) activity data along with associated metadata, including peptide sequences, target bacterium, strain information, and experimental conditions (Filippov and Shkirskaya, 2019). GRAMPA combines data from five major AMP repositories— DBAASP (Pirtskhalava et al., 2016), APD (Wang et al., 2016), DADP (Novković et al., 2012), DRAMP (Fan et al., 2016), and YADAMP (Piotto et al., 2012) resulting in 6,760 unique peptide sequences and 51,345 antimicrobial activity measurements. Among the bacterial targets, *Escherichia coli* constitutes the largest subset with 9,150 measurements, followed by *Staphylococcus aureus* with 8,954 measurements. Given the higher sample availability for *E. coli* and consistency with the evaluation protocol of the reference study MSCMamba (He et al., 2025), our experiments were conducted exclusively on the *E. coli* subset of the GRAMPA dataset. To assess the generalizability of the proposed approach, we additionally report results on the Staphylococcus aureus and Pseudomonas aeruginosa subset as secondary evaluation datasets.

#### Unlabeled AMP dataset

A large collection of unlabeled antimicrobial peptide sequences was assembled from multiple publicly available AMP sources. The sequences were merged, cleaned, and filtered to remove redundant or low-quality entries. To further reduce redundancy and enhance sequence diversity, clustering was performed at a 60% sequence-identity threshold, resulting in 47,215 retained peptide sequences. This dataset was subsequently used for semi-supervised model training to complement the experimentally labeled MIC dataset.

#### Escherichia coli dataset

The original 9,150 *E. coli* samples were standardized by removing sequences containing non-standard amino acids and excluding entries with invalid MIC values. Duplicate peptide records were consolidated using the geometric mean of MIC measurements, resulting in 4,793 unique peptides for model evaluation (Supplementary Section S1).

#### Staphylococcus aureus dataset

Applying the same preprocessing pipeline, the original 8,954 records were filtered accordingly, yielding a final dataset of 4,381 sequences (Supplementary Section S2).

#### Pseudomonas aeruginosa dataset

The same pipeline was also applied to the *P. aeruginosa* dataset, reducing the original 4,966 records to 2,722 curated sequences.

#### Stratified Data Splits

The final dataset was partitioned into training, validation, and test sets using randomized stratification of MIC values, which were discretized into five quantile-based bins to maintain distributional balance. Protein language embeddings and physicochemical descriptors were extracted prior to subset allocation for model training and evaluation (Supplementary Section S3).

Figure 1 summarizes the feature extraction framework. Sequence-derived features included embedding representations, physicochemical descriptors, BLOSUM62-based features, and positional encodings.

**Figure 1.**
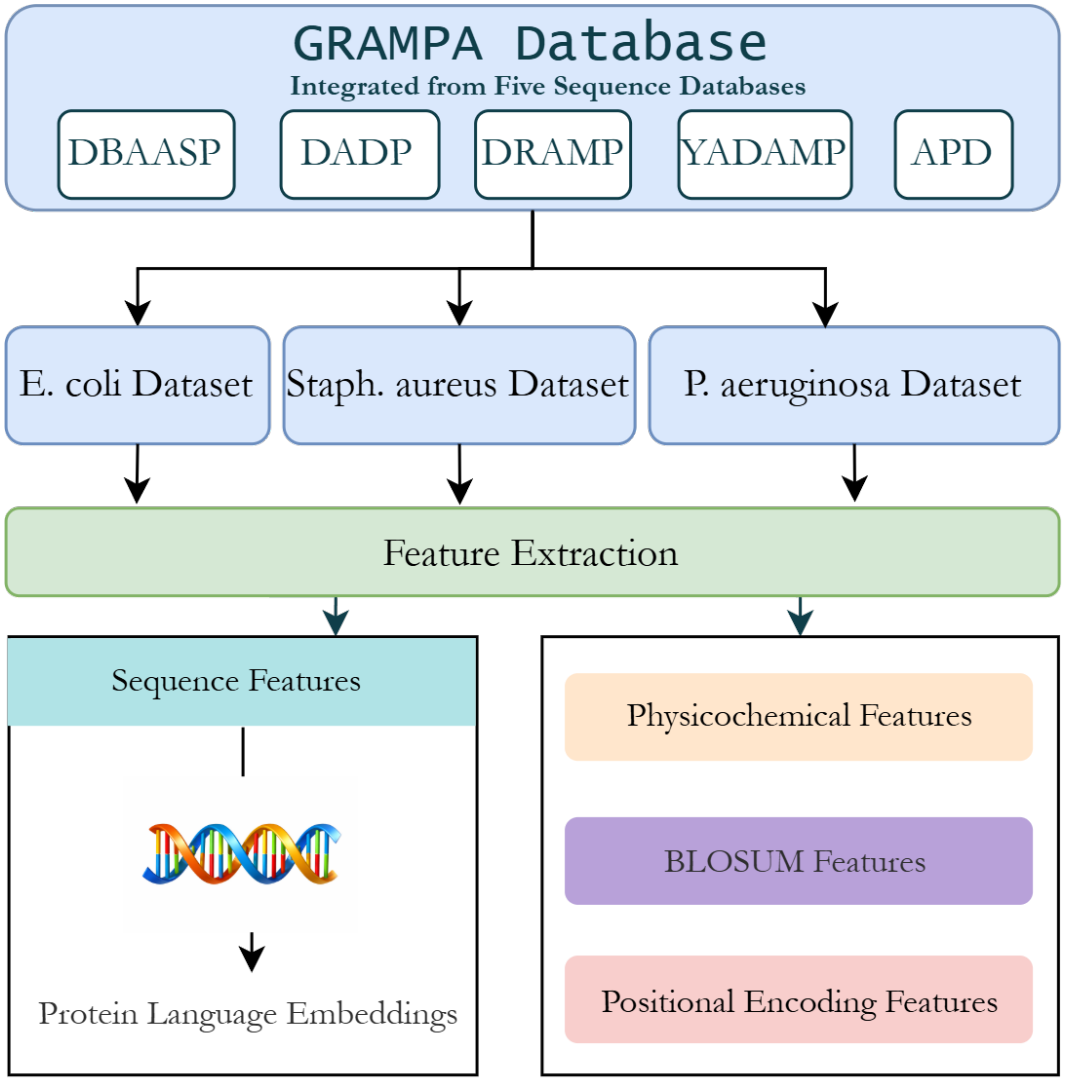
Workflow for deriving model-ready features from the GRAMPA database, which provides peptide sequences, antimicrobial activity values, and associated metadata. Model-ready representations are generated by extracting sequence embeddings and engineered features, including physicochemical descriptors, BLOSUM62-based features, and positional encodings.

### Model Architecture

We propose a trust-weighted gated modulation protein language model (PLM) framework, *TW-GMProt*, for the prediction of antimicrobial activity (Fig. 2). The architecture (Fig. 2A) consists of three complementary branches: (a) a ProtT5-based branch that captures contextual sequence representations, (b) a CNN-based branch that extracts local motif patterns and modulates the protein language model representation through gated interaction, and (c) an engineered sequence branch that integrates physicochemical descriptors, BLOSUM62 features, and sinusoidal positional encodings. The outputs of the ProtT5 and CNN branches are combined through gated modulation to refine the contextual protein representation. In parallel, engineered sequence inputs are projected into a compact representation through a multilayer perceptron. Next, the output features from gated modulation are combined with the engineered feature representation through concatenation fusion to predict MIC.

**Figure 2.**
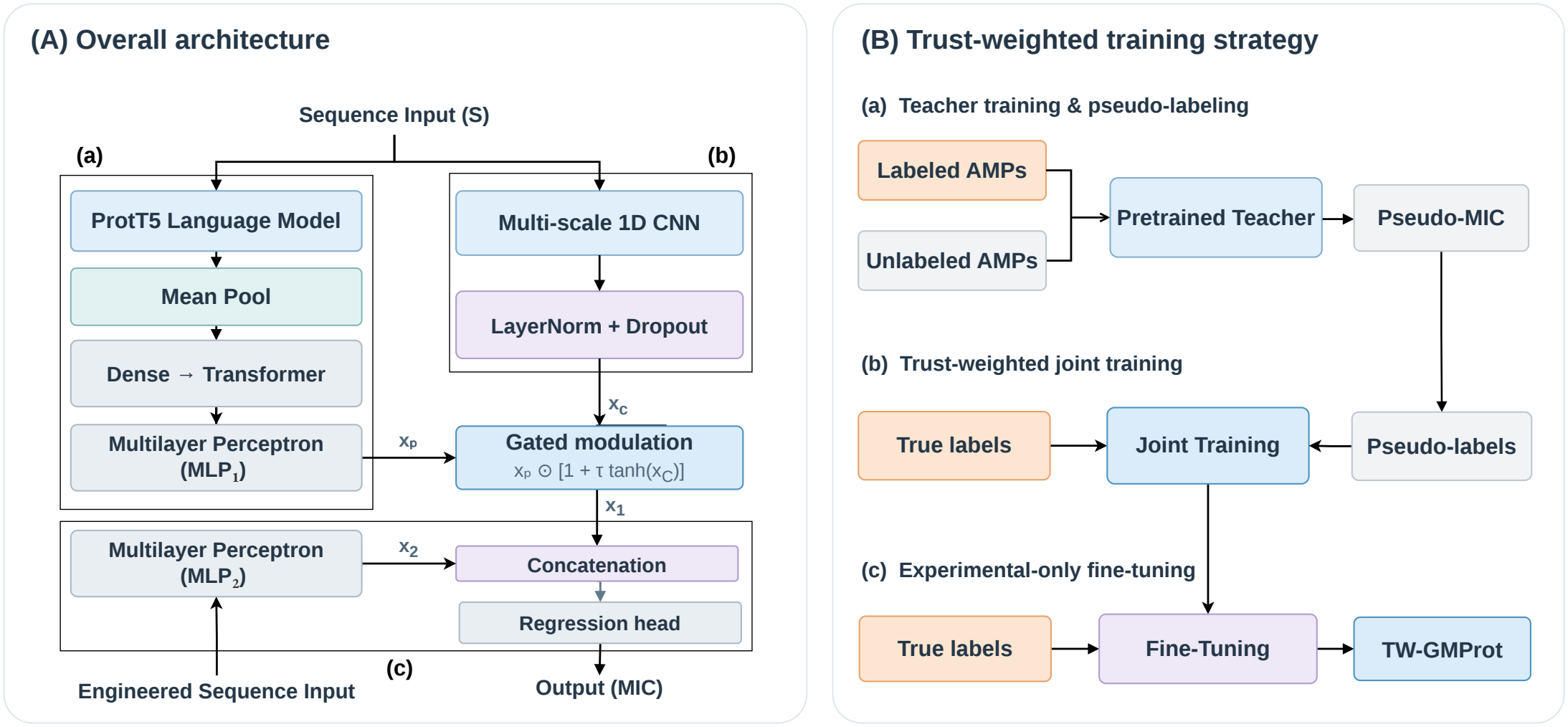
Overview of the proposed TW-GMProt framework. (A) Overall architecture: ProtT5-derived sequence representations are refined through CNN-guided gated modulation and integrated with complementary engineered sequence features via additive fusion for MIC prediction. (B) Trust-weighted training strategy: Once trained on experimentally measured MIC values, the model serves as a teacher to generate pseudo-MIC labels for a large collection of unlabeled AMP sequences (B(a)). Subsequently, experimental and pseudo-labeled samples are jointly used to retrain the model, with full weight assigned to experimental labels and reduced trust weights assigned to pseudo-labels (B(b)). The jointly trained model is then fine-tuned exclusively on experimentally labeled samples to obtain the final model (B(c)).

The limited availability of experimental activity labels often hinders reliable model performance in data-scarce settings, motivating the exploration of semi-supervised approaches. To address this limitation, we adopted a trust-weighted semi-supervised learning strategy (Fig. 2B), in which an initial model trained on experimental MIC measurements serves as a teacher to generate pseudo-MIC labels for a substantially larger collection of unlabeled antimicrobial peptide sequences. The trust-weighted strategy assigns full weight to experimentally measured MIC labels while incorporating pseudo-labeled samples with reduced weights during joint training. However, the resulting intermediate model may be susceptible to biases introduced by imperfect pseudo-labels. To mitigate these potential biases, the jointly trained model is subsequently fine-tuned exclusively on experimentally labeled samples, yielding the final TW-GMProt model.

### Gated Modulation Protein Language Embedding

Sequence embeddings are extracted using the pretrained protein language model ProtT5 (Fig. 2A(a)). For each residue, a 1024-dimensional embedding is generated, resulting in residue-level representations. ProtT5 was selected as the primary sequence feature extractor because its large-scale self-supervised pretraining on diverse protein sequences enables it to generate rich contextual representations that encode sequence-dependent biochemical and biophysical information. Previous studies have also demonstrated the effectiveness of ProtT5 embeddings for representing peptide sequences and various protein-level properties. Although alternative protein language models, including ESM-based models (Lin et al., 2023; Rao et al., 2021) and ProtBERT, also provide informative contextual representations, their relative effectiveness can vary across downstream tasks and sequence-length distributions. In our setting, ProtT5 provides a suitable representation for short peptide sequences by capturing residue-level context and sequence patterns without requiring explicit evolutionary information. Therefore, ProtT5 was adopted as the primary sequence representation model.

Let 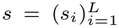 denote a protein sequence of length *L*, where *s*_*i*_ *∈* Ω represents the *i*-th amino acid from alphabet Ω. The pretrained ProtT5 encoder generates contextual residue embeddings ProtT5(*s*) *∈* ℝ^*L×*1024^, which are globally pooled to obtain a fixed-length sequence representation *z*_0_ *∈* ℝ^1024^. This representation is projected to 512 dimensions via a dense layer Dense_1_, followed by a 10-head Transformer encoder (TF) to capture task-specific residue interactions and higher-order dependencies. The resulting features are further refined using a multilayer perceptron (MLP_1_), consisting of two fully connected layers (256 *→* 128), yielding the final sequence embedding *x*_*P*_ *∈* ℝ^128^:

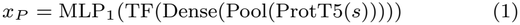

In addition, CNN-based embedding features (Fig. 2A(b)) were derived to capture local motif patterns that may not be fully represented by ProtT5 embedding. While ProtT5 embeddings effectively capture long-range dependencies, they may overlook important local sequence patterns, including motifs, residue spacing, periodicity, and short-range compositional features. Functional activity measures, such as minimum inhibitory concentration (MIC), are not determined solely by long-range contextual information. To address this limitation, we incorporated a CNN-based branch to guide and refine the sequence representation. Input sequences are first embedded and processed through four parallel 1D convolutional layers (kernel sizes 3, 5, 7, and 11) to capture multi-scale patterns, with padded positions masked to prevent spurious contributions. The resulting feature maps are global-max-pooled, concatenated, and projected into a 128-dimensional sequence embedding, followed by layer normalization and dropout for regularization, yielding the output feature *x*_*C*_ *∈* ℝ^128^:

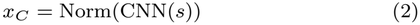

While CNN-based architectures effectively capture local sequence motifs, they are prone to overfitting when trained on limited peptide datasets. Therefore, instead of using the CNN branch as an independent feature extractor, we employ it as a gated refinement module for ProtT5 embeddings. Only a small fraction of CNN-derived signals (*τ* = 0.2%) is allowed to modulate the ProtT5 representation, enabling the recovery of complementary local motif information that may be underrepresented in global language model embeddings; this threshold was chosen based on ablation studies, as lower values failed to capture sufficient local motif information, while higher values introduced noise.

The gated modulation is implemented through a tanh-based residual gating mechanism that generates a bounded mask in the range [–1, 1] from CNN-derived representations. The bounded residual design stabilizes feature magnitudes while enabling controlled, bidirectional adjustment of pretrained embeddings without disrupting their underlying semantic structure. By integrating local motif cues with global contextual embeddings in a controlled manner, the model produces a more robust and predictive sequence representation *x*_1_:

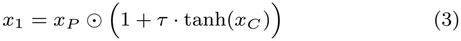

where *τ* denotes the gating coefficient controlling the modulation ratio, and *⊙* represents Hadamard product.

### Engineered Sequence Features

Physicochemical properties serve as important predictor variables for guiding the model in determining antimicrobial activity, as they capture the underlying mechanisms by which compounds interact with bacterial membranes and other molecular targets. We extracted a total of 32 physicochemical descriptors *x*^phys^ *∈* ℝ^32^ using modLamp (Müller et al., 2017). These include 10 global descriptors, 20 amino acid composition features, and two Eisenberg scale descriptors. The selection of these features was guided by two key considerations. First, the feature set should sufficiently capture the most significant physicochemical properties that play a crucial role in antimicrobial activity. Second, care was taken to avoid redundant or highly overlapping descriptors, thereby reducing the risk of model overfitting. Feature selection was further validated through systematic ablation studies. A detailed list of physicochemical features is provided in Supplementary Section S4.

BLOSUM62 features capture residue similarities based on evolutionary substitutions, aiding the model in recognizing functionally conserved patterns (VERCAUTEREN et al., 2024). While BLOSUM45 is designed for distantly related sequences and BLOSUM80 for highly conserved sequences, BLOSUM62 offers a balanced scoring scheme suitable for the moderately similar sequences typical of short AMPs (Henikoff and Henikoff, 1992; Eddy, 2004). For each peptide, we extracted a 20-dimensional BLOSUM62 composition vector *x*^blosum^ *∈* ℝ^20^ by averaging the corresponding rows of the substitution matrix across all residues in the sequence:

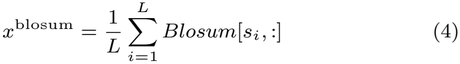

This provides a fixed-length sequence-level representation that captures evolutionary and functional similarity between residues.

The biological activity of a peptide is governed by the precise sequence of its amino acids. To capture these order-specific features, we computed positional encodings by using the sinusoidal encoding from transformers (Vaswani et al., 2017), providing position-aware representations that help recognize spatial arrangements. Each sequence was encoded as a matrix of shape (*d, D* = 32) where *d* is the sequence length. Since sequence lengths vary, the encodings were padded to a maximum length of 128. The sinusoidal positional encoding *PE* for position *p* and dimension index *i* is given by:

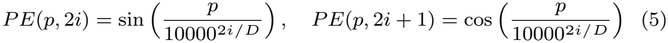

Global pooling over positions produces a single vector *x*^pos^ *∈* ℝ^32^ for each peptide:

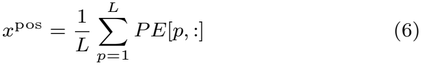

Physicochemical (32), BLOSUM (20), and positional encoding (32) features were concatenated to form an 84-dimensional engineered sequence feature vector *x*_eng_ *∈* ℝ^84^, capturing global, evolutionary, and order-based information:

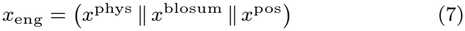

### Prediction of MIC

The engineered sequence representation *x*_eng_ is projected through a multilayer perceptron (MLP_2_) consisting of two fully connected layers, yielding a 128-dimensional descriptor representation *x*_2_:

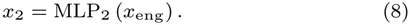

The gated protein-language-model representation *x*_1_ and engineered sequence representation *x*_2_ are integrated through concatenation fusion. The resulting multimodal representation is passed through a final dense regression layer to estimate MIC:

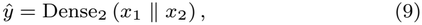

where *ŷ* denotes the predicted MIC value.

### Trust-Weighted Pseudo-Label Learning

Experimentally determined MIC measurements are substantially more reliable than computationally inferred activity values; however, experimentally annotated peptide datasets remain limited in size. To exploit the substantially larger pool of unlabeled peptide sequences, we adopted trust-weighted pseudo-label learning strategy.

Let 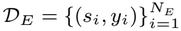 denote the experimentally labeled dataset, where *y*_*i*_ represents the measured MIC of peptide *s*_*i*_. An initially trained model 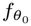 is used as a teacher model to infer MIC values for a larger unlabeled peptide collection 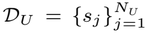. For each unlabeled peptide, a pseudo-MIC value is generated as

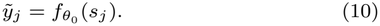

The resulting pseudo-labeled dataset is therefore defined as

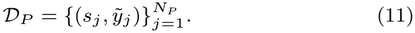

Because pseudo-labels originate from model inference rather than experimental measurement, they are not treated as equivalent to ground-truth MIC values. Instead, each experimental sample is assigned full trust, 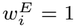, whereas pseudo-labeled samples receive substantially smaller weights. To prevent the much larger pseudo-labeled dataset from dominating optimization purely because of sample count, the base pseudo-label weight is normalized according to the relative dataset sizes:

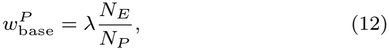

where *λ* controls the total contribution of pseudo-labeled data relative to experimentally labeled data. Then, the joint weighted training regression objective becomes

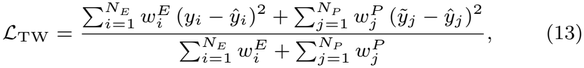

where 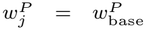 in the basic trust-weighted formulation.

This weighting strategy allows pseudo-labeled peptides to expand the sequence space observed during optimization while ensuring that experimentally measured MIC values remain the dominant supervision source.

### Two-Stage Training

Initially, the *model* is trained exclusively on experimentally labeled dataset to establish a teacher model (Fig. 2A), which is subsequently used to generate pseudo-MIC labels for unlabeled peptide sequences. The model then undergoes two sequential training stages to obtain the final *m*odel (Fig. 2B).

In the first stage, the model is trained jointly on experimentally labeled and pseudo-labeled samples to obtain an intermediate model (Fig. 2B(b)). In the second stage, this intermediate model is fine-tuned exclusively on experimentally labeled samples to mitigate potential biases introduced by imperfect pseudo-labels (Fig. 2B(c)).

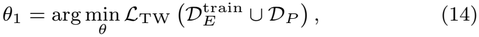

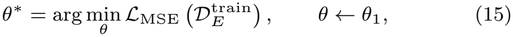

where *θ*_1_ denotes the parameters learned from joint trust-weighted training and *θ*^*\**^ denotes the final parameters obtained after experimental-label-only fine-tuning.

### Implementation Details

Training was performed end-to-end by minimizing mean squared error (MSE) loss and implemented using TensorFlow 2.13/Python 3.10 on an NVIDIA A100 GPU. ProtT5 embeddings were extracted using ‘Rostlab/prot t5 xl uniref50’ from Hugging Face Transformers (4.57.3) with PyTorch 2.9.1.

## Results

We evaluated the proposed framework for antimicrobial activity prediction against *Escherichia coli* using five-fold stratified cross-validation. Model performance was assessed using RMSE, Kendall’s *τ*, *R*^2^, and Pearson’s *r*.

### Overall Performance

The proposed TW-GMProt model was benchmarked against eight established methods: MSCMamba (He et al., 2025), IAMPCN (Xu et al., 2023), AI4AMP (Lin et al., 2021), AMPDLMD (Cao et al., 2023), KNN (Witten and Witten, 2019), AMPpred-MFA (Li et al., 2023), TranslmbAMP (Pang et al., 2022), and WittenCNN (Witten and Witten, 2019). TW-GMProt outperformed all competing methods, achieving an RMSE of **0.435**, Kendall’s *τ* of **0.583**, *R*^2^ of **0.623**, and Pearson’s *r* of **0.793** (Fig. 3, Table 1). To assess robustness to random initialization, we conducted 50 runs across 10 random seeds and five independent hold-out datasets. Paired Wilcoxon signed-rank tests on seed-specific RMSE values indicated a statistically significant improvement for TW-GMProt over MSCMamba (*p* = 1.95 *×* 10^*−*3^).

**Table 1.** Performance comparison on AMP activity prediction.

| **Method** | **RMSE ↓** | **Kendall ↑** | ***R*<sup>2</sup> ↑** | **Pearson *r* ↑** |
|---|---|---|---|---|
| IAMPCN | 0.580 | 0.395 | 0.347 | 0.596 |
| WittenCNN | 0.546 | 0.454 | 0.422 | 0.664 |
| AI4AMP | 0.582 | 0.410 | 0.343 | 0.599 |
| AMPDLMD | 0.607 | 0.377 | 0.285 | 0.583 |
| K-NN | 0.563 | 0.424 | 0.385 | 0.645 |
| AMPpred-MFA | 0.591 | 0.385 | 0.323 | 0.601 |
| TransImbAMP | 0.573 | 0.428 | 0.363 | 0.649 |
| MSCMamba | 0.524 | 0.462 | 0.467 | 0.684 |
| **TW-GMProt** | **0.435**±0.011 | **0.583**±0.012 | **0.623**±0.016 | **0.793**±0.011 |

**Figure 3.**
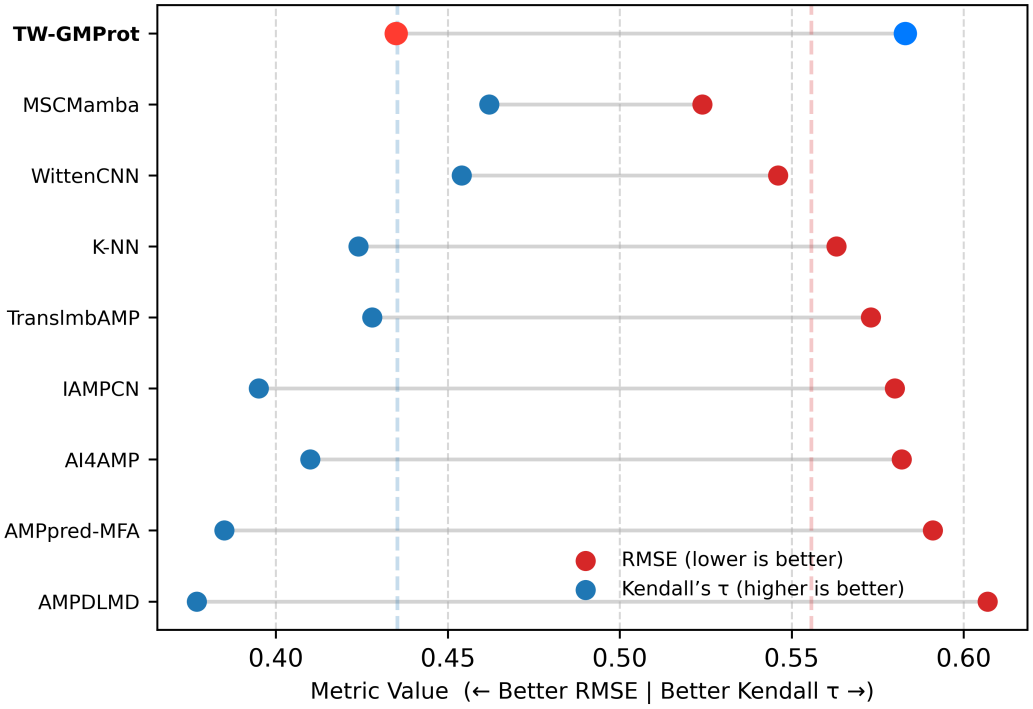
Performance comparison of eight baseline methods, and TW-GMProt based on RMSE and Kendall’s *τ*.

To assess the contribution of the trust-weighted semi-supervised learning strategy, we compared the model trained exclusively on experimentally measured MIC-labeled data with its semi-supervised counterpart incorporating additional pseudo-labeled samples. The semi-supervised approach consistently outperformed the experimental-only baseline, achieving a 9.56% reduction in RMSE (0.481 to 0.435) and relative improvements of 17.30% in Kendall’s *τ* (0.497 to 0.583), 27.40% in *R*^2^ (0.489 to 0.623), and 12.64% in Pearson’s *r* (0.704 to 0.793). These findings highlight the effectiveness of incorporating trust-weighted pseudo-labeled data in improving quantitative antimicrobial activity prediction, as summarized in Table 2.

**Table 2.** Performance comparison of experimental-only and trust-weighted semi-supervised training strategies on the *E*.*Coli* dataset, highlighting the contribution of pseudo-labeled data.

| **Strategy** | **RMSE** ↓ | **Kendall** τ ↑ | *R*<sup>2</sup> ↑ | **Pearson** *r* ↑ |
|---|---|---|---|---|
| Experimental-only | 0.481±0.010 | 0.497±0.011 | 0.489±0.017 | 0.704±0.012 |
| Semi-supervised | **0.435**±0.011 | **0.583**±0.012 | **0.623**±0.016 | **0.793**±0.011 |

### Ablation Studies

We systematically evaluated the performance of various components of TW-GMProt for MIC prediction. The ablation study was conducted exclusively on the original experimental dataset, without incorporating any semi-supervised data. Sequence-derived architectures (Table 3) integrated ProtT5 embeddings with Transformer and CNN modules. Ablation studies were designed to assess the contribution of each architectural component and feature modality.

**Table 3.**
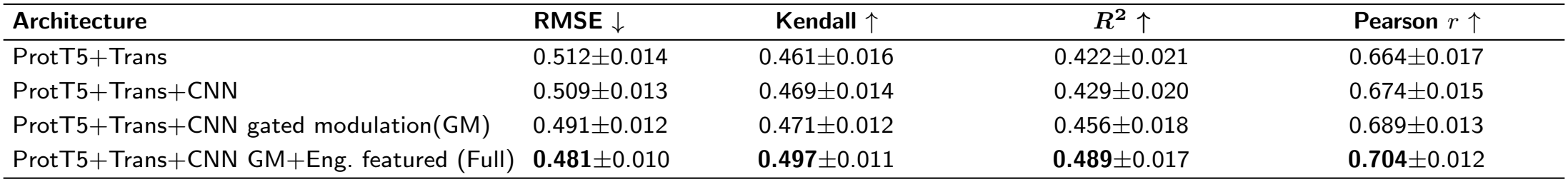
Ablation analysis of different sequence-derived architectures for the model.

Progressive refinement of sequence-derived models, starting from a ProtT5+Transformer baseline to CNN and CNN-gated variants (Table 3), consistently improved all evaluation metrics. This highlights the importance of local motif extraction and adaptive feature reweighting in antimicrobial activity prediction. The highest performance was achieved by the CNN gated modulation variant, following the integration of engineered features.

Different variants of the model were also evaluated to assess the impact of weighted attention and cross-attention (Table 4) as alternatives to concatenation. The attention-weighted fusion approach consistently reduced performance across all evaluation metrics. Although cross-attention achieved performance comparable to direct feature concatenation, it did not yield measurable improvements and resulted in lower *R*^2^ values. This may be because applying cross-attention to already learned high-level representations increases model complexity without introducing substantial new information, potentially leading to overfitting or the incorporation of redundant features. These findings suggest that the learned sequence representations are already sufficiently informative for activity prediction, and that more sophisticated fusion mechanisms provide limited benefit while potentially introducing noise.

**Table 4.**
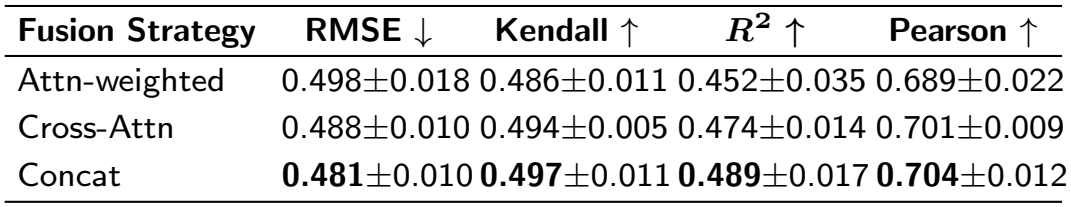
Performance ablation of the model utilizing attention-weighted, cross-attention, and concatenation-based feature fusion strategies.

Additionally, the model was evaluated using embeddings from ProtT5, ProtBert, and ESM2 (Table 5). ProtT5 and ProtBert produce 1024-dimensional residue embeddings, while ESM2 outputs 1280-dimensional features. ProtT5 achieved the best performance, likely because its span-masked objective better captures global semantics and long-range inter-residue dependencies. In contrast, ProtBERT’s token-level masking provides less contextual information, while ESM2 may produce noisier representations for short, flexible peptides due to its optimization for large, structured proteins.

**Table 5.**
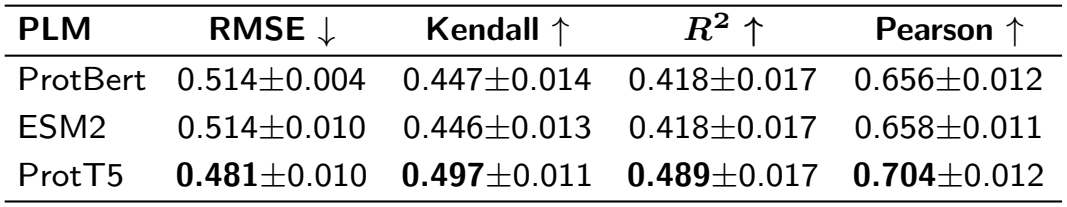
Ablation with different Protein Language Models.

Overall, the ablation studies demonstrate that each component of TW-GMProt contributes to improved MIC prediction, with CNN-based modulation and engineered sequence features providing the considerable performance gains. More complex feature fusion strategies offered no additional benefit over simple feature concatenation.

### Model Generalizability across Bacterial Species

To assess cross-species applicability, we further trained and evaluated the model on smaller experimental datasets of *Pseudomonas aeruginosa* and *Staphylococcus aureus*, without incorporating pseudo-labels. The model achieved comparable predictive performance on the corresponding independent test sets. Although benchmark results for these datasets are unavailable, the consistent performance highlights the potential of the proposed framework for antimicrobial activity prediction against diverse antimicrobial species (Table 6).

**Table 6.** Cross-species evaluation on the *Staphylococcus aureus (S.A.)* and *Pseudomonas aeruginosa (P.A.)* datasets.

| **Dataset** | **Split %** | **RMSE ↓** | **Kendall ↑** | ***R*<sup>2</sup> ↑** | **Pearson ↑** |
|---|---|---|---|---|---|
| *P.A.* | 70/15 | 0.505±0.025 | 0.482±0.020 | 0.406±0.031 | 0.651±0.021 |
| *P.A.* | 80/10 | 0.479±0.024 | 0.488±0.066 | 0.434±0.044 | 0.672±0.034 |
| *S.A.* | 70/15 | 0.491±0.010 | 0.492±0.030 | 0.488±0.020 | 0.707±0.020 |
| *S.A.* | 80/10 | 0.488±0.011 | 0.496±0.024 | 0.494±0.016 | 0.712±0.024 |

### Interpretability Analysis

To understand the significance of different input features for MIC prediction, we performed interpretability analyses using CNN gating ablation and Integrated Gradients (Sundararajan et al., 2017). The CNN gating contribution analysis revealed a subtle yet meaningful modulation across peptides. As seen in figure 4A, by comparing predictions from the sequence model with and without CNN gating (mean = −0.009), we observed that the CNN branch fine-tunes sequence-derived embeddings without dominating predictions.

**Figure 4.**
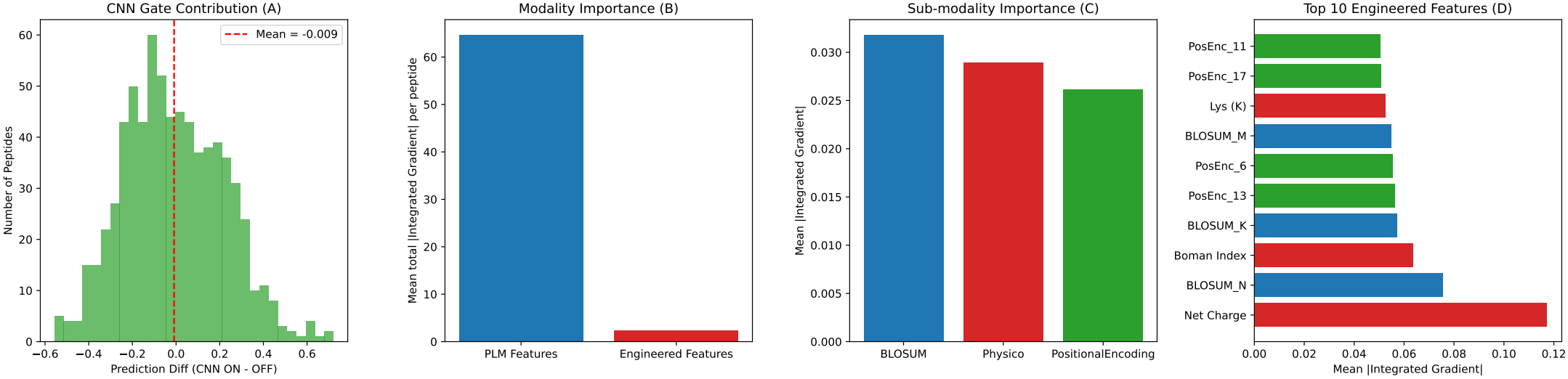
Model interpretability and feature contribution analysis: (A) CNN gating subtly modulates sequence embeddings. (B) PLM features dominate MIC prediction, while engineered features provide valuable complementary signals. (C) Sub-modality importance highlights contributions from engineered sequence categories. (D) Top ten engineered sequence features from the total 84 descriptors.

At the modality level, attribution analysis was performed by computing the mean absolute Integrated Gradients (|IG|) for each feature. The results indicate that ProtT5-based protein language model embeddings contribute substantially more to MIC prediction, while engineered sequence features provide complementary information that further supports predictive performance (Fig. 4B). Sub-modality importance analysis further reveals that BLOSUM features are the highest contributors, followed by physicochemical and positional encoding features (Fig. 4C). Figure 4D presents the top ten key drivers among the 84 engineered sequence features.

### Model Assessment for Discovering AMP-Associated Motifs

We evaluated 256 peptide sequences from the *E. coli* test dataset to assess whether the model captures antimicrobial peptide (AMP)-associated sequence patterns. A motif discovery strategy inspired by TF-MoDISco (Shrikumar et al. (2018)) was adopted to interpret model predictions.

Integrated Gradients (IG) were computed for each peptide sequence to derive residue-level attribution scores from the CNN-based sequence encoder, while keeping the remaining model inputs fixed. Each peptide sequence was represented using learned embeddings, and IG values were obtained by linearly interpolating between a baseline constructed from the model’s learned padding token (PAD) embedding and the actual sequence embeddings over 64 steps. Gradients of the model output with respect to the interpolated embeddings were integrated to obtain residue-level attribution scores, quantifying the contribution of individual amino acids to the model prediction relative to the selected baseline.

To identify localized regions of high attribution, IG scores were converted into short subsequences (seqlets) using an adaptive peak-based segmentation strategy. Residues exceeding the 85th percentile of attribution scores were selected as seed positions. From each seed, contiguous regions were expanded bidirectionally until the attribution signal fell below a lower expansion threshold (65th percentile) or the maximum seqlet length constraint (25 residues) was reached. A minimum seqlet length of three residues was imposed. Overlapping segments were excluded, and proximal segments were merged using a maximum gap of one residue. This adaptive strategy reduces dependence on predefined fixed-length windows and facilitates the identification of localized attribution patterns.

The extracted seqlets were subsequently clustered using a hybrid similarity metric combining *k*-mer sequence similarity and IG-derived similarity. Sequence similarity (*k* = 3) was assigned a higher weight (*α* = 0.90), while IG similarity provided complementary attribution information. Agglomerative hierarchical clustering was performed using the resulting distance matrix and a similarity threshold of 0.70. Motif consensus sequences were then generated from the resulting clusters.

To enhance biological interpretability, motif clusters were filtered using sequence-complexity and physicochemical criteria associated with antimicrobial peptides. Candidate motifs were assessed based on net charge, mean hydrophobicity, a helical hydrophobic-moment approximation, and a composite AMP validity score. Clusters were retained if they satisfied the minimum consensus-strength criterion (0.40), contained at least two seqlets, exhibited a nominal net positive charge of at least +1, and achieved an AMP validity score greater than 1.5. A sequence-complexity filter was also applied to exclude selected repetitive or compositionally biased patterns. The retained clusters were ranked using a motif importance score combining mean absolute IG attribution, the square root of seqlet support, and consensus strength, with attribution magnitude scaled relative to the maximum absolute seqlet attribution.

Following filtering and ranking, 11 candidate AMP-associated motifs were identified, comprising 24 supporting seqlets from 21 unique peptide sequences. The highest-ranked motifs were LIK, FLGALFK, LGK, FIK, and KIT, with motif importance scores of 0.850, 0.744, 0.638, 0.580, and 0.520, respectively (Fig. 5B; Table 7). Several motifs exhibited recurring combinations of hydrophobic residues, particularly leucine (L), isoleucine (I), and phenylalanine (F), together with positively charged lysine (K). The extended motif FLGALFK contained multiple hydrophobic residues and a terminal lysine, whereas shorter motifs such as LIK, FIK, and IVK displayed localized hydrophobic–cationic sequence patterns. Such residue combinations are consistent with compositional characteristics frequently observed in antimicrobial peptides, where cationic and hydrophobic residues can contribute to membrane association. However, their direct involvement in membrane binding or disruption cannot be established from sequence attribution alone.

**Table 7.** Ranked AMP-associated motifs identified through IG-based motif discovery. Importance denotes the attribution-based ranking score, whereas AMP score represents the physicochemical validity heuristic. Hydrophobicity is the mean Kyte–Doolittle value.

| Rank | Motif | Support | Importance | Charge | Hydro. | AMP<br>Score |
|---|---|---|---|---|---|---|
| 1 | **LIK** | 2 | **0.850** | +1 | 1.47 | 4.65 |
| 2 | **FLGALFK** | 2 | **0.744** | +1 | 1.53 | 4.17 |
| 3 | **LGK** | 2 | **0.638** | +1 | -0.17 | 3.27 |
| 4 | FIK | 2 | 0.580 | +1 | 1.13 | 4.65 |
| 5 | KIT | 2 | 0.520 | +1 | -0.03 | 3.54 |
| 6 | LKA | 2 | 0.504 | +1 | 0.57 | 4.35 |
| 7 | IVK | 3 | 0.488 | +1 | 1.60 | 4.63 |
| 8 | AAAAAAKL | 3 | 0.295 | +1 | 1.34 | 3.84 |
| 9 | KFP | 2 | 0.197 | +1 | -0.90 | 2.01 |
| 10 | KTF | 2 | 0.136 | +1 | -0.60 | 2.47 |
| 11 | VIR | 2 | 0.128 | +1 | 1.40 | 4.66 |

**Figure 5.**
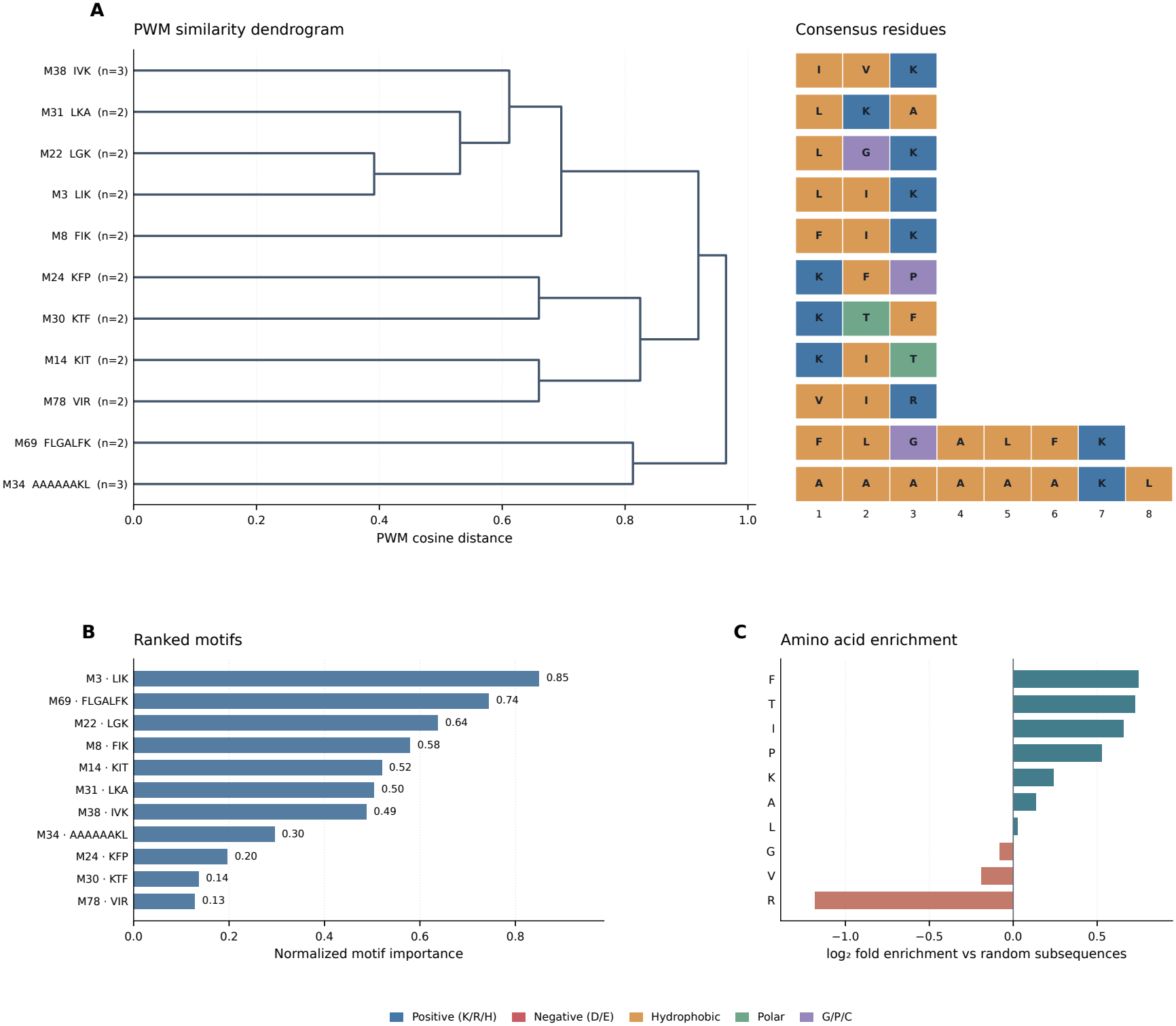
Integrated Gradients (IG)-based identification and characterization of candidate AMP-associated sequence motifs. (A) Hierarchical organization of 11 retained motifs using position-weight matrix (PWM) cosine distance, accompanied by a consensus-residue heatmap. Motifs were initially identified through seqlet clustering using a hybrid similarity metric combining *k*-mer sequence similarity (*k* = 3, *α* = 0.90) and IG attribution similarity. Dendrogram branch heights represent PWM cosine distance, while leaf labels indicate consensus sequences and supporting seqlet counts. Residues in the heatmap are colored according to their physicochemical categories. (B) Ranking of candidate motifs based on attribution magnitude, seqlet support, and consensus strength. The highest-ranked motifs include LIK, FLGALFK, LGK, FIK, and KIT, which exhibit recurring hydrophobic and cationic residue combinations. (C) Amino acid enrichment expressed as log_2_ fold enrichment relative to randomly sampled peptide subsequences matched to the observed seqlet-length distribution. Positive and negative values indicate relative enrichment and depletion, respectively. Residues with fewer than two observed occurrences were excluded from visualization.

The identified motifs ranged from three to eight residues in length and exhibited a nominal net charge of +1 under the adopted residue-based charge approximation. Their mean Kyte–Doolittle hydrophobicity values varied considerably, reflecting differences in residue composition. For example, LIK and FLGALFK showed positive mean hydrophobicity values of 1.47 and 1.53, respectively, whereas LGK exhibited a slightly negative value of *−*0.17 (Table 7). These findings indicate that the retained motifs possess heterogeneous physicochemical characteristics rather than a uniform hydrophobic profile. Although several motifs showed relatively high calculated hydrophobic moments, these values were derived under an idealized helical assumption and do not independently establish amphipathic secondary structures.

Furthermore, amino acid enrichment analysis was performed by comparing the residue composition of the retained motif seqlets with 2,000 randomly sampled peptide subsequences matched to the observed seqlet-length distribution. The analysis revealed increased representation of phenylalanine (F; 1.68-fold), threonine (T; 1.65-fold), isoleucine (I; 1.58-fold), and lysine (K; 1.18-fold) relative to the random background (Fig. 5C; Supplementary Table S3). In contrast, arginine (R) showed reduced representation (0.44-fold), while leucine (L) displayed approximately comparable frequencies between the observed and background subsequences (1.02-fold). The enrichment of phenylalanine and isoleucine suggests preferential representation of specific hydrophobic residues within the selected motifs, whereas the modest enrichment of lysine indicates the presence of cationic sequence characteristics. Nevertheless, these compositional patterns should be interpreted cautiously because the motif-selection procedure explicitly incorporates a positive-charge criterion, and several enriched residues occur at low frequencies.

Collectively, the motif analysis identified recurring hydrophobic– cationic sequence patterns associated with the model’s MIC predictions, suggesting that the learned representations capture sequence characteristics compatible with known AMP-associated physicochemical properties. However, most motif clusters were supported by only two or three seqlets, and motif importance was calculated from absolute IG attribution, which does not distinguish contributions toward increased or decreased predicted MIC. Therefore, the identified patterns should be regarded as exploratory model-associated motifs rather than experimentally validated antimicrobial determinants.

## Discussion and Conclusion

We presented TW-GMProt, a semi-supervised gated modulation framework for predicting the minimum inhibitory concentration (MIC) of antimicrobial peptides. The model integrates protein language model embeddings with CNN-guided local sequence refinement and biophysical priors. Our ablation experiments demonstrated that gated sequence modulation and multimodal feature integration improve MIC prediction performance. To address the performance limitations arising from the limited availability of experimental data, we adopted a trust-weighted semi-supervised learning strategy.

Our model trained exclusively on experimental data outperforms all evaluated benchmarks, while the semi-supervised approach achieves further higher improvements (*p* = 1.95 *×* 10^*−*3^) in predictive performance. These results highlight two key contributions of our work: the effectiveness of the proposed model architecture and the successful integration of pseudo-labels for the antimicrobial activity prediction.

Integrated Gradients analysis indicated substantial contributions from physicochemical and BLOSUM62-derived features, while positional encodings also contributed to peptide ranking. These findings highlight the complementary roles of physicochemical, evolutionary, and sequence-order information alongside contextual protein representations. The motif discovery analysis suggests that the model captures sequence characteristics commonly associated with antimicrobial activity. The identified motifs exhibited recurring combinations of cationic and hydrophobic residues, consistent with known physicochemical properties of antimicrobial peptides. These findings provide supporting evidence that the learned representations capture biologically plausible sequence patterns potentially relevant to peptide–membrane interactions.

Although our semi-supervised strategy improves the utilization of limited experimental data, pseudo-labels may propagate systematic prediction errors if their contributions are not appropriately controlled. The trust-weighted formulation mitigates this risk by reducing the influence of pseudo-labeled samples and subsequently fine-tuning the model exclusively on experimentally labeled data. Future work could further refine this strategy by incorporating peptide-specific confidence estimates, such as ensemble disagreement or uncertainty-aware weighting, rather than assigning a uniform trust level to all pseudo-labeled samples.

While the present study focused on antimicrobial peptide activity, the proposed framework is not inherently restricted to a single species and could potentially be extended to other peptide activity prediction tasks, including antiviral Singh Bist et al. (2023); Bist et al. (2024, 2026) and antifungal applications. Further evaluation across diverse organisms and independent datasets is needed to establish the broader generalizability of the proposed approach.

In conclusion, this study introduced *TW-GMProt*, a trust-weighted gated modulation framework for peptide activity prediction that leverages semi-supervised learning to integrate pseudo-labeled peptide data with experimentally measured samples. The proposed framework offers a promising approach for improving activity prediction and peptide ranking, potentially facilitating candidate prioritization for experimental validation and accelerating therapeutic peptide discovery.

## Data Availability and Implementation

The datasets and trained models supporting the findings of this work are archived on Zenodo at https://doi.org/10.5281/zenodo.20740473. The source code is available at https://github.com/SCSE-Biomedical-Computing-Group/GMProt

## Supplementary Material

Supplementary materials are available online.

## References

M. Alzain, H. Daghistani, T. Shamrani, Y. Almoghrabi, Y. Daghistani, O. S. Alharbi, A. M. Sait, M. Mufrrih, W. Alhazmi, M. A. Alqarni, et al. Antimicrobial peptides: Mechanisms, applications, and therapeutic potential. Infection and Drug Resistance, pages 4385–4426, 2025.

B. Behera, R. Singh, K. Sharma, A. Rai, S. Singh, and B. Balan. A comprehensive review of advanced strategies to combat antimicrobial resistance. Archives of Microbiology, 207(11): 281, 2025.

S. Bhattarai, K. T. Chong, and H. Tayara. Gan-ml: Advancing anticancer peptide prediction through innovative deep convolution generative adversarial network data augmentation technique. Chemometrics and Intelligent Laboratory Systems, 262:105390, 2025.

P. S. Bist, H. Tayara, and K. T. Chong. Generative ai in the advancement of viral therapeutics for predicting and targeting immune-evasive sars-cov-2 mutations. IEEE Journal of Biomedical and Health Informatics, 2024.

S. Bist, K. T. Chong, and H. Tayara. Unveiling viral escape mechanisms with machine learning: A transformative approach to mutation analysis for sars-cov-2 and beyond. IEEE Transactions on Computational Biology and Bioinformatics, 2026.

Cao, C. Ge, X. Wang, P. J. Harvey, Z. Zhang, Y. Ma, X. Wang, X. Jia, M. Mobli, D. J. Craik, et al. Designing antimicrobial peptides using deep learning and molecular dynamic simulations. Briefings in Bioinformatics, 24(2): bbad058, 2023.

Z. Chen, C. Ji, W. Xu, J. Gao, J. Huang, H. Xu, G. Qian, and J. Huang. Uniamp: enhancing amp prediction using deep neural networks with inferred information of peptides. BMC bioinformatics, 26(1):10, 2025.

S. R. Eddy. Where did the blosum62 alignment score matrix come from? Nature biotechnology, 22(8):1035–1036, 2004.

L. Fan, J. Sun, M. Zhou, J. Zhou, X. Lao, H. Zheng, and H. Xu. DRAMP: A comprehensive data repository of antimicrobial peptides. Scientific Reports, 6:1–7, 2016. ISSN 20452322. doi: 10.1038/srep24482. URL http://dx.doi.org/10.1038/srep24482.

A. N. Filippov and S. A. Shkirskaya. Deep learning regression model for antimicrobial peptide design. Membrany i Membrannye Tekhnologii, 9(5):325–333, 2019. doi: 10.1134/s221811721905002x.

M. He, Y. Jiang, Y. Yang, K. Gong, X. Jiang, and Y. Tian. MSCMamba: Prediction of Antimicrobial Peptide Activity Values by Fusing Multiscale Convolution with Mamba Module. Journal of Physical Chemistry B, 129(7):1956–1965, 2025. ISSN 15205207. doi: 10.1021/acs.jpcb.4c07752.

S. Henikoff and J. G. Henikoff. Amino acid substitution matrices from protein blocks. Proceedings of the National Academy of Sciences, 89(22):10915–10919, 1992.

C. Li, Q. Zou, C. Jia, and J. Zheng. Amppred-mfa: an interpretable antimicrobial peptide predictor with a stacking architecture, multiple features, and multihead attention. Journal of Chemical Information and Modeling, 64(7):2393–2404, 2023.

T.-T. Lin, L.-Y. Yang, I.-H. Lu, W.-C. Cheng, Z.-R. Hsu, S.-H. Chen, and C.-Y. Lin. Ai4amp: an antimicrobial peptide predictor using physicochemical property-based encoding method and deep learning. Msystems, 6(6):e00299–21, 2021.

Z. Lin, H. Akin, R. Rao, B. Hie, Z. Zhu, W. Lu, N. Smetanin, R. Verkuil, O. Kabeli, Y. Shmueli, et al. Evolutionary-scale prediction of atomic-level protein structure with a language model. Science, 379(6637):1123–1130, 2023.

K. H. Min, K. H. Kim, M.-R. Ki, and S. P. Pack. Antimicrobial peptides and their biomedical applications: A review. Antibiotics, 13(9):794, 2024.

A. T. Müller, G. Gabernet, J. A. Hiss, and G. Schneider. modlAMP: Python for antimicrobial peptides. Bioinformatics (Oxford, England), 33(17):2753–2755, 2017. ISSN 13674811. doi: 10.1093/bioinformatics/btx285.

M. Novković, J. Simunić, V. Bojović, A. Tossi, and D. Juretić. DADP: The database of anuran defense peptides. Bioinformatics, 28(10):1406–1407, 2012. ISSN 13674803. doi: 10.1093/bioinformatics/bts141.

Y. Pang, L. Yao, J. Xu, Z. Wang, and T.-Y. Lee. Integrating transformer and imbalanced multi-label learning to identify antimicrobial peptides and their functional activities. Bioinformatics, 38(24):5368–5374, 2022.

S. P. Piotto, L. Sessa, S. Concilio, and P. Iannelli. YADAMP: Yet another database of antimicrobial peptides. International Journal of Antimicrobial Agents, 39(4):346–351, 2012. ISSN 09248579. doi: 10.1016/j.ijantimicag.2011.12.003. URL http://dx.doi.org/10.1016/j.ijantimicag.2011.12.003.

M. Pirtskhalava, A. Gabrielian, P. Cruz, H. L. Griggs, R. B. Squires, D. E. Hurt, M. Grigolava, M. Chubinidze, G. Gogoladze, B. Vishnepolsky, V. Alekseev, A. Rosenthal, and M. Tartakovsky. DBAASP v.2: An enhanced database of structure and antimicrobial/cytotoxic activity of natural and synthetic peptides. Nucleic Acids Research, 44(D1):D1104– D1112, 2016. ISSN 13624962. doi: 10.1093/nar/gkv1174.

R. M. Rao, J. Liu, R. Verkuil, J. Meier, J. Canny, P. Abbeel, T. Sercu, and A. Rives. Msa transformer. In International conference on machine learning, pages 8844–8856. PMLR, 2021.

A. Shrikumar, K. Tian, Ž. Avsec, A. Shcherbina, A. Banerjee, M. Sharmin, S. Nair, and A. Kundaje. Technical note on transcription factor motif discovery from importance scores (tfmodisco) version 0.5. 6.5. arXiv preprint arXiv:1811.00416, 2018.

P. Singh Bist, H. Tayara, and K. To Chong. Sars-escape network for escape prediction of sars-cov-2. Bioinformatics, 24(3):bbad140, 2023. Briefings in

M. Sundararajan, A. Taly, and Q. Yan. Axiomatic attribution for deep networks. In International conference on machine learning, pages 3319–3328. PMLR, 2017.

A. Vaswani, N. Shazeer, N. Parmar, J. Uszkoreit, L. Jones, A. N. Gomez, L. Kaiser, and I. Polosukhin. Attention is all you need. In Advances in Neural Information Processing Systems, volume 30, 2017.

K. Vercauteren, P. Meysman, and K. Laukens. Tcrblosum: An amino acid substitution matrix for sensitive alignment of distant epitope-specific tcrs. BRIEFINGS IN BIOINFORMATICS, 26(1), 2024.

G. Wang, X. Li, and Z. Wang. APD3: The antimicrobial peptide database as a tool for research and education. Nucleic Acids Research, 44(D1):D1087–D1093, 2016. ISSN 13624962. doi: 10.1093/nar/gkv1278.

Y. Wang, V. Liang, N. Yin, S. Liu, and E. Segal. Sgac: A graph neural network framework for imbalanced and structure-aware amp classification. arXiv preprint arXiv:2412.16276, 2024.

J. Witten and Z. Witten. Deep learning regression model for antimicrobial peptide design. BioRxiv, page 692681, 2019.

W. Xing, J. Zhang, C. Li, Y. Huo, and G. Dong. iampattenpred: a novel antimicrobial peptide predictor based on bert feature extraction method and cnn-bilstm-attention combination model. Briefings in bioinformatics, 25(1), 2023.

J. Xu, F. Li, C. Li, X. Guo, C. Landersdorfer, H.-H. Shen, A. Y. Peleg, J. Li, S. Imoto, J. Yao, et al. iampcn: a deep-learning approach for identifying antimicrobial peptides and their functional activities. Briefings in Bioinformatics, 24(4): bbad240, 2023.

Q.-Y. Zhang, Z.-B. Yan, Y.-M. Meng, X.-Y. Hong, G. Shao, J.-J. Ma, X.-R. Cheng, J. Liu, J. Kang, and C.-Y. Fu. Antimicrobial peptides: mechanism of action, activity and clinical potential. Military Medical Research, 8(1):48, 2021.

