## Supplementary Information for "TW-GMProt: Trust-Weighted Semi-Supervised Gated Modulation Protein Language Model for Antimicrobial Peptide Activity Prediction"

#### S1 Construction of the *Escherichia coli* Database

To ensure data standardization, we preprocessed the original 9,150 *E. coli* samples. Samples containing non-standard amino acids (B, J, O, U, and X) were excluded. As multiple minimum inhibitory concentration (MIC) measurements may exist for the same peptide due to integration from different databases, duplicate records were consolidated by computing the geometric mean of the MIC values.

Furthermore, experimental entries with negative MIC values were removed to ensure data validity and consistency, following the processing strategy adopted in MSCMamba (He et al., 2025). After excluding 874 invalid MIC records, we identified 1,800 duplicate peptide groups, comprising 5,363 duplicate samples. As a result, a curated *E. coli* dataset consisting of 4,793 unique peptides was constructed and used for network evaluation.

### S2 Overview of Datasets

**Table S1:** Dataset construction statistics and source database distributions.

| Dataset | Count | Remarks |
|---|---|---|
| **Reference dataset** | | |
| GRAMPA | 51,345 | Activity measurements from 6,760 unique peptide sequences aggregated across multiple AMP databases |
| ***E. coli*** **dataset** | | |
| Original | 9,150 | Subset extracted from GRAMPA |
| Final | 4,793 | Curated after filtering, deduplication, and MIC consolidation |
| DBAASP | 3,760 | Final dataset records by source database |
| APD | 478 | |
| DADP | 256 | |
| DRAMP | 160 | |
| YADAMP | 139 | |
| ***Staphylococcus aureus*** **dataset** | | |
| Original | 8,954 | Subset extracted from GRAMPA |
| Final | 4,381 | Curated after filtering, deduplication, and MIC consolidation |
| DBAASP | 3,325 | Final dataset records by source database |
| APD | 470 | |
| DADP | 285 | |
| DRAMP | 157 | |
| YADAMP | 144 | |
| ***Pseudomonas aeruginosa*** **dataset** | | |
| Original | 4,966 | Subset extracted from GRAMPA |
| Final | 2,722 | Curated after filtering, deduplication, and MIC consolidation |

### S3 Generation of Five Stratified Data Splits

The final *E. coli* dataset was split into training (3,259), validation (815), and test (719) sets by using independent randomized stratification based on MIC values. This splitting strategy follows the protocol used in MSCMamba (He et al., 2025) to ensure a comparable evaluation framework. Similarly, the *Staphylococcus aureus* and *Pseudomonas aeruginosa* datasets were split into train/validation/test sets at 70:15:15 and 80:10:10. Since MSC-Mamba provides no benchmarks for this dataset, we used these standard splits to assess generalization. MIC values were discretized into five quantile-based bins to preserve their distribution across splits. ProtT5 (Elnaggar et al., 2022) sequence embeddings and Mod-Lamp (Müller et al., 2017) physicochemical descriptors were extracted from the complete datasets and subsequently assigned to the corresponding training, validation, and test subsets for model training and evaluation.

#### S4 Physicochemical features

**Table S2:** Physicochemical features ued for antimicrobial peptide activity prediction.

| Feature Name | Description and Role in Activity Prediction |
|---|---|
| **Global descriptors (10)** | |
| Molecular Weight (MW) | Total molecular mass of the peptide; affects diffusion, penetration, and stability |
| Length | Number of amino acid residues; correlates with mechanism of action and membrane insertion |
| Net Charge | Total charge at physiological pH; electrostatic attraction to bacterial membranes |
| Charge Density | Net charge normalized by peptide length; concentration of positive charge for efficient activity |
| Isoelectric Point (pI) | pH at which net charge is zero; predicts cationic behavior under physiological pH |
| Aliphatic Index | Relative volume occupied by aliphatic side chains; hydrophobic interactions with lipid membranes |
| Aromaticity | Fraction of aromatic residues; anchoring and insertion into membranes |
| Instability Index | Indicator of peptide stability in vitro; predicts peptide stability and functional persistence |
| Boman Index | Protein-binding potential; affects selectivity |
| Hydrophobic Ratio | Fraction of hydrophobic residues; drives amphipathicity and membrane permeabilization |
| **Amino acid composition (20)** | |
| Frequency of each amino acid (A, C, D, E, F, G, H, I, K, L, M, N, P, Q, R, S, T, V, W, Y) | Quantify enrichment/depletion of specific residues; encodes general sequence composition trends; captures variable-length sequence properties |
| **Eisenberg scale (2)** | |
| Eisenberg Hydrophobic Moment | Measure of amphipathicity |
| Eisenberg Global Hydrophobicity | Average hydrophobicity over sequence |

**Table S3:** Amino acid enrichment analysis of IG-derived AMP-associated motifs relative to randomly sampled peptide subsequences. Fold enrichment greater than 1 indicates relative enrichment, whereas values below 1 indicate depletion. Frequencies include pseudocount smoothing.

| AA | Motif Count | Random Count | Motif Freq. | Random Freq. | Fold Enrich. | $\log_2$ Enrich. |
|---|---|---|---|---|---|---|
| F | 10 | 463 | 0.108 | 0.065 | **1.678** | **0.747** |
| T | 4 | 201 | 0.046 | 0.028 | **1.654** | **0.726** |
| I | 11 | 540 | 0.119 | 0.075 | **1.576** | **0.656** |
| P | 2 | 128 | 0.026 | 0.018 | 1.441 | 0.527 |
| K | 20 | 1284 | 0.211 | 0.179 | 1.182 | 0.242 |
| A | 18 | 1247 | 0.191 | 0.174 | 1.099 | 0.136 |
| L | 11 | 835 | 0.119 | 0.116 | 1.020 | 0.028 |
| G | 4 | 352 | 0.046 | 0.049 | 0.946 | -0.080 |
| E | 0 | 39 | 0.005 | 0.005 | 0.938 | -0.093 |
| V | 5 | 464 | 0.057 | 0.065 | 0.877 | -0.189 |
| M | 0 | 45 | 0.005 | 0.006 | 0.814 | -0.297 |
| Y | 0 | 48 | 0.005 | 0.007 | 0.764 | -0.389 |
| R | 2 | 420 | 0.026 | 0.059 | 0.440 | -1.183 |
| D | 0 | 92 | 0.005 | 0.013 | 0.400 | -1.320 |
| Q | 0 | 100 | 0.005 | 0.014 | 0.369 | -1.440 |
| N | 0 | 103 | 0.005 | 0.014 | 0.358 | -1.482 |
| C | 0 | 124 | 0.005 | 0.017 | 0.298 | -1.749 |
| H | 0 | 171 | 0.005 | 0.024 | 0.216 | -2.211 |
| S | 0 | 223 | 0.005 | 0.031 | 0.166 | -2.593 |
| W | 0 | 297 | 0.005 | 0.041 | 0.125 | -3.006 |

#### References

A. Elnaggar, M. Heinzinger, C. Dallago, G. Rehawi, Y. Wang, L. Jones, T. Gibbs, T. Feher, C. Angerer, M. Steinegger, D. Bhowmik, and B. Rost. ProtTrans: Toward Understanding the Language of Life Through Self-Supervised Learning. *IEEE Transactions on Pattern Analysis and Machine Intelligence*, 44(10):7112–7127, 2022. ISSN 19393539. doi: 10.1109/TPAMI.2021.3095381.

M. He, Y. Jiang, Y. Yang, K. Gong, X. Jiang, and Y. Tian. MSCMamba: Prediction of Antimicrobial Peptide Activity Values by Fusing Multiscale Convolution with Mamba Module. *Journal of Physical Chemistry B*, 129(7):1956–1965, 2025. ISSN 15205207. doi: 10.1021/acs.jpcb.4c07752.

A. T. Müller, G. Gabernet, J. A. Hiss, and G. Schneider. modlAMP: Python for antimicrobial peptides. *Bioinformatics (Oxford, England)*, 33(17):2753–2755, 2017. ISSN 13674811. doi: 10.1093/bioinformatics/btx285.
